# The selective culture and enrichment of major rumen bacteria on three distinct anaerobic culture media

**DOI:** 10.1101/2024.08.23.608987

**Authors:** Alice M. Buckner, Laura Glendinning, Juan M. Palma Hidalgo, Jolanda M. van Munster, Mark Stevens, Mick Watson, C. Jamie Newbold

## Abstract

Ruminants play an important part in global food security, but also emit methane which contributes to global warming. Microbes in the rumen strongly influence the energy retention efficiency from the host’s plant-based diet and produce methane as a by-product. While thousands of novel microbial genomes have been assembled from metagenome sequence data, their culturability is ill-defined. Here different media were used to isolate microbes from rumen fluid. 34 genera were grown, and the majority belonged to the phylum *Bacillota* (75.28% ± 6.34), *Bacteroidota* (19.99% ± 4.85), *Pseudomonadota* (2.46% ± 2.01), and *Actinomycetota* (2.09% ± 1.07). The most abundant genera were *Selenomonas* (28.08% ± 11.71), *Streptococcus* (22.67% ± 6.06), *Prevotella* (18.71% ± 4.02), and unclassified *Lachnospiraceae* (11.50% ± 2.54). When comparing the mean relative abundance of these genera between media, 31 were significantly enriched on at least one medium. The composition of the source rumen fluid was vastly different to those cultured. *Bacteroidota* (52.53% ± 5.10) predominated, with by *Bacillota* (41.00% ± 3.96), the archaeal *Euryarchaeota* (5.12% ± 1.94), *Pseudomonadota* (1.22% ± 0.78), and *Actinomycetota* (0.12% ± 0.08) comprising the rest. The most abundant genera were *Prevotella* (29.13% ± 4.16), *Butyrivibrio* (18.21% ± 2.08), *Succiniclasticum* (15.57% ± 5.03), unclassified *Bacteroidetes* (13.91% ± 1.67), and unclassified *Prevotellaceae* (9.50% ± 2.01). These data further emphasise the importance of using defined media to selectively enrich for different microbial taxa. This is essential to understand the complex workings of the rumen microbes to enhance digestion efficiency and reduce the loss of energy as methane.

## 2 Introduction

The majority of ruminant digestion occurs in the forestomach termed the reticulorumen, referred to as the rumen in this paper, which contains a community of microorganisms that aid the host in its digestion of plant matter (1). This microbial community consists of a number of taxa that are common across a wide variety of ruminants, the majority of which are bacteria (2,3). However, the type and abundance of other taxa vary by factors such as species, breed, diet, or location (4). This composition has a great influence on the efficiency of ruminant digestion (5,6). Around 18.4% of global greenhouse gas emissions are produced by the agricultural industry, with 5.8% of global emissions coming from livestock and manure alone (7), indicating the need to mitigate this and optimise the efficiency of ruminant digestion (2).

Studying the rumen microbiome to find methods to ensure it is as efficient as possible is key to this goal, with traditional research methods involving culturing the organisms *in vitro* (8,9). However, rumen microbes are fastidious and require strict anaerobic techniques to isolate them (10). Following the development of Next Generation Sequencing (NGS) techniques, the use of metagenomics and metabarcoding strategies to analyse the community composition has become increasingly popular due to the wealth of information they can provide (11). There are advantages and disadvantages to each of these methods (12,13), however it has been recognised that the combination of sequencing and culturing can provide a more complete view of the rumen microbiome (10,14,15).

Members of the rumen microbiome currently isolated in culture do not represent the full diversity and complexity of the community, and taxonomic classification of those currently in culture is as yet incomplete (16–18), with many rumen microbes likely to have been misclassified (16). The fact that some microbes seem to resist cultivation indicates a gap in knowledge of their basic metabolism and interactions with their environment (10). The phyla *Bacillota* (heterotypic synonym; *Firmicutes)*, *Pseudomonadota* (homotypic synonym; *Proteobacteria)*, and *Actinomycetota* (homotypic synonym; *Actinobacteria)* are found most often in culture, with *Bacteroidota* (homotypic synonym; *Bacteroidetes)* being under represented (16). Being able to culture these microbes *in vitro* is key to understanding how to optimise ruminant digestion, however the estimated proportion of culturable microbes in the rumen varies, spanning from < 1% (19) to 23% (15). Methods to increase this proportion include using a higher number of technical replicates, multiple media, and multiple dilutions of rumen fluid (15,20). This study aims to; i) culture a range of dominant, fast-growing rumen bacteria, ii) determine if any of these bacteria are selectively enriched on a range of well-established, widely used media for the culture of rumen microbes, and iii) to compare the range of bacteria grown here to the whole community of bacteria in the original rumen fluid. This will provide a better understanding of which rumen microbes can be cultured on each of these commonly used media, and how abundant the microbes grown are in the rumen community.

## 3 Methods

### 3.1 Sample collection and microbial culture

Samples of rumen fluid, for the purpose of inoculation and culturing, were collected from four fistulated, female, Jersey cows, aged 5 to 12 years, grazing a sward of perennial ryegrass. Rumen fluid was collected via the fistula, squeezed into a pre-warmed thermos, and transported back to the laboratory for inoculation within two hours of collection. The samples were squeezed through two layers of muslin into a CO_2_ gassed container and kept in a water bath set to 39°C. This was then serially diluted in anaerobic diluent to achieve a 10^-2^ dilution of the original rumen fluid. The anaerobic diluent was based on that from Bryant and Burkey (1952) (21), however the concentration of cysteine was reduced to match that present in the media (0.51 g/L). In addition to the dilution series, 8 mL of the strained rumen fluid was spun at 3018 *g* for 20 minutes before the supernatant was removed and the pellet stored at −80°C.

McSweeney et al. (2005) describe five types of anaerobic culture media commonly used for isolation of rumen bacteria and archaea (19). Based on a pilot experiment (data not shown) three media were chosen based on their ability to support microbial growth (based on the level of change in optical density over time after inoculation with rumen fluid). These three media were composed of two non-selective media, one containing rumen fluid (Med2 (22)) and the other not (Med10 (23)), and one selective medium (MedTC (19)) which contained rumen fluid as well as a range of other substrates not present in the non-selective media. The non-selective media aimed to mimic the entire rumen environment to culture a wide range of rumen microbes, while the selective medium contained additional substrates to support additional microbes, such as ones that are xylanolytic, pectinolytic, or that rely on the presence of trace elements.

All the stock solutions that were used to make up the media were made in aerobic conditions. The anaerobic diluent and three types of media were made using these component solutions under anaerobic conditions using 100% CO_2_ gas. The water was added to a heat-proof flask and put on a hot plate with a magnetic stirrer and temperature probe. The hot plate was set to 95°C and stirrer to 300 rpm. The liquid and solid media ingredients were added, and when the mixture reached 95°C the CO_2_ line was added and the hot plate set to 55°C. When the mixture reached 55°C the reducing agents were added. When the mixture had no red tint, the medium was dispensed into 10 mL Hungate tubes for the media or 100 mL Wheaton tubes for the anaerobic diluent. The tube was gassed with CO_2_, the anaerobic solution added, and a cap and bung quickly used to seal the tube. The tubes were then autoclaved at standard conditions, allowed to cool to room temperature, and the caps sterilised with ethanol before the heat unstable ingredients were added using a needle and syringe. An overview of the components of the media and anaerobic diluent is given in Supplementary Table 4.

For each cow and each medium, four controls were processed and incubated alongside their respective samples. Two of the controls consisted of sterile media only, while two were mock inoculated with 1 mL of sterile anaerobic diluent. Ethanol was used to sterilise the caps of the tubes of media, and 1 mL of the 10^-2^ dilution was added to five replicate tubes of media. The optical density (590 nm) of the cultures was measured just after inoculation (time 0), after 24 hours, and after 48 hours of incubation at 39°C. After the 48-hour measurement 8 mL of each sample was removed and spun at 4,500 rpm for 20 minutes. The supernatant was discarded and the pellet stored at −80°C for DNA extraction. 2 mL of sterile 25% glycerol was added to the remaining 2 mL of culture, and was stored at −80°C.

### 3.2 Determining microbial composition

The pellets of the original rumen fluid sample and the cultures (along with negative and reagent-only controls) were subject to DNA extraction, 16S amplification, and massively-parallel sequencing before bioinformatics was used to determine the OTUs present. A microbial standard community (ZymoBIOMICS Microbial Community Standard) was also used to determine how effective and accurate these steps were. The relevant results are described in Supplementary Figure 1.

DNA from the pellet was extracted using a modified protocol based on the Qiagen DNeasy® PowerLyzer® PowerSoil® Kit and Qiagen QIAamp® Fast DNA Stool Mini Kits. The majority of the protocol followed the QIAamp® Fast DNA Stool Mini Kit protocol, with an additional glass bead beating step on centrifuged culture samples with the supernatant removed (as described in DNeasy® PowerLyzer® PowerSoil® with modifications described in Supplementary Note 1), and eluting the DNA in a lower volume (50 µL). DNA was extracted from the Zymo microbial standard with InhibitX added to 75ul of the standard before the protocol was carried out as per the other samples.

16S rDNA amplification was then carried out on the eluted DNA. 12 µL of Q5 Hot Start High-Fidelity 2X Master Mix, 9 µL of nuclease free water, 1.25 µL of the forward primer (10nM), and 1.25 µL of the reverse primer (10nM) was added to either 1 μl or 0.2 μl of the template DNA, depending on which provided the highest quantity of amplicon product. Barcoded primers (5′–TATGGTAATTGTGTGCCAGCMGCCGCGGTAA–3′ and 5′–AGTCAGTCAGCCGGACTACHVGGGTWTCTAAT–3′) were used to target the V4 region before being cycled through a PCR machine with a heated lid, running at 95°C for 2 minutes followed by 30 cycles of 95°C for 20 seconds, 55°C for 15 seconds, 72°C for 5 minutes followed by 72°C for 10 minutes. The product was then purified using Ampure magnetic bead purification in a 1:1 ratio, quantified using Nanodrop and Qubit, and visualised using gel electrophoresis. Using the HS Qubit values for the concentration of amplified DNA in each sample, the DNA was combined to form a pool with equimolar concentrations of DNA from each sample.

Sequencing of the single pool was carried out by Edinburgh Genomics using Illumina MiSeq V2 chemistry, with 500 cycles producing an output of around 11 million 300 base pair paired-end reads. As very few reads were produced for three of the four rumen fluid samples, and due to the importance of having these samples in the dataset, they were sequenced again in another identical Miseq run and then substituted into the initial dataset.

Bioinformatic analysis was carried out using Mothur v. 1.48.0., and follows the Miseq SOP (https://mothur.org/wiki/miseq_sop/) (24). Steps in the sections ‘Reducing sequencing and PCR errors’ and ‘Processing improved sequences’ were followed exactly, with the exception of Archaea not being removed in the remove.lineage command. OTUs were generated using the commands given in the ‘OTU’ section of the ‘Preparing for analysis’ section (without the use of cluster.split), before consensus sequences for each OTU were generated using the classify.otu command given in the ‘Phylotypes’ section. Databases used were Version 138.1 of the SILVA database (https://mothur.org/wiki/silva_reference_files/) with the ‘full-length sequences and taxonomy references’ modified to contain just the V4 region, and Version 18 of the ‘16S rRNA reference (RDP)’ database (https://mothur.org/wiki/rdp_reference_files/).

R v. 4.2.2. was used with the package Phyloseq v. 1.42.0. (25) to collate the metadata, OTU table, and taxonomy table, and the protocol followed was based on that as recommended by the package developer and others including (26–28). The OTU table, taxonomy table, metadata, and tree were imported from MOTHUR into R. They were combined using the Phyloseq package to give 8,152 taxa by 6 taxonomic ranks from 96 samples by 9 sample variables.

Sampling coverage was calculated to determine which samples to remove by calculating the number of reads per sample (Supplementary Figure 1). From the histogram, many samples had very few reads, with the scatterplot showing a large cluster of samples at the bottom at around 1e+02 with the rest scattered around 1e+05. This is also echoed in the line graph showing many low abundance samples followed by significant step up to around 3,000 reads. The Good’s coverage was then calculated, which quantifies the relationship between the number of sequences and the number of OTUs only present once in each sample, with 7 of the 60 samples having a Good’s coverage of < 99%. Due to this, samples with fewer than (or equal to) 3,075 reads or 92.3% Good’s coverage were removed to give 8,129 taxa across 82 samples.

Sampling depth was calculated to determine which low abundance OTUs to remove by looking at the total number of reads per OTU across all samples. The total number of reads was 5,256,392. If any OTUs had a relative abundance of less than 0.1% across all cultures, they were removed. These OTUs are unlikely to be able to be reliably cultured again as, for bacteria with a relative abundance of < 0.1%, over 1000 tubes of media would have to be inoculated for these bacteria and archaea to be cultured once. This resulted in a dataset containing 43 taxa from 82 samples. Finally, any OTUs present in the cultures or controls, but not present in any of the rumen fluid samples, were removed, giving 34 taxa. Despite this step removing 8,118 OTUs, only 11.7% of the total reads were discarded.

To determine if there was a difference in the OTUs found in the medium-only controls between the three media, the α-diversity (Observed, Shannon, and Chao1 calculated using the Microbiome R package) and the β-diversity (weighted UniFrac) were calculated. An ANOVA was carried out to determine if there was a significant difference between the α-diversity metrics by medium, with a post-hoc Tukey test used if p < 0.05 to investigate pairwise interactions. For the β-diversity a PERMANOVA, using adonis2 from the Vegan package, was used to determine if there was a significant difference between the β-diversity by medium, with a post-hoc pairwise PERMAVOVA run if p < 0.05. Following this, each medium-only control was compared to the cultures grown on this medium to determine if the OTU diversity had changed. A two-sample T Test was carried out to determine the difference between the α-diversities of the medium-only controls and the cultures grown on that medium, and a PERMANOVA was used for the β-diversity. Next, the α-diversity metrics and β-diversity was compared between the cultures by medium, with an ANOVA (and post-hoc Tukey) and PERMANOVA (and post-hoc pairwise PERMANOVA with manual Bonferroni adjustment for multiple testing) used as described above. Finally, the difference in OTU relative abundance was determined using the same method above using a Kruskal–Wallis test (along with a post-hoc Dunn test if p < 0.05). The 8 most abundance OTUs across all cultures were shown in a boxplot as they make up over 80% of the dataset.

## 4 Results

### 4.1 Identification of operational taxonomic units (OTUs) present in rumen fluid and cultured on different media

Rumen fluid, diluted 1/100 in anaerobic diluent, was added to three distinct types of anaerobic liquid media (Med2, Med10, and MedTC) and incubated at 39°C for 48 hours. After the rumen fluid had been cultured, the microbes incubated, and their DNA extracted; their 16S V4 regions were amplified using unique barcoded primers, pooled, and sequenced. The raw sequencing data reads were then processed to generate operational taxonomic units (OTUs) which were classified to genus level. Of the 8,152 OTUs generated, the majority were filtered out of the dataset, either due to there being a low number of reads for that OTU ( < 0.1% abundance across all samples) or a low number of reads for a particular sample ( < 3075 reads or a Good’s Coverage of < 92.3%) (Supplementary Figure 1). At the end of the filtering steps, and after removing any OTUs not present in the original rumen fluid, 34 OTUs were present in the cultures. The 8,118 OTUs removed only accounted for 11.7% of the entire number of reads in the dataset.

### 4.2 Microbial DNA in the basal medium

The three media chosen to cultivate the rumen microbes were based on those described by McSweeney et al. (2005) ^19^. The media Med2 and MedTC contained sterile clarified rumen fluid as a medium constituent, whereas the medium Med10 did not. After sequencing the DNA extracted from all the samples, including medium-only controls, it was determined that there was rumen microbe DNA present in the medium from this sterile clarified rumen fluid. However there was a minimal effect on the presence of this DNA on the results (Supplementary Figure 2). On further comparison between each medium and its respective cultures, there were sufficient significant differences between the OTUs in the basal medium and those in the inoculated cultures to indicate the selective growth of different rumen microbes (Supplementary Figure 3, Supplementary Figure 4, Supplementary Figure 5).

### 4.3 Microbes grown in different media

After 48 hours it was observed, based on the optical density, that the community of microbes in the culture tubes inoculated with fresh rumen fluid as a whole had grown through the lag and exponential phases of growth (data not shown), and had reached the stationary phase. At this stage, the community composition was determined. For all the relative abundances below, the variation metric given is the standard deviation. Of the 34 taxa found across all the cultured samples, *Bacillota* (75.28% ± 6.34) was the most common phylum cultured, based on their relative abundance, followed by *Bacteroidota* (19.99% ± 4.85), *Pseudomonadota* (2.46% ± 2.01), and *Actinomycetota* (2.09% ± 1.07) (Figure 1a). The most common genera cultured across all media, based on their relative abundance, were *Selenomonas* (28.08% ± 11.71), *Streptococcus* (22.67% ± 6.06), *Prevotella* (18.71% ± 4.02), and unclassified *Lachnospiraceae* (11.50% ± 2.54) (Figure 1b).

**Figure 1.**
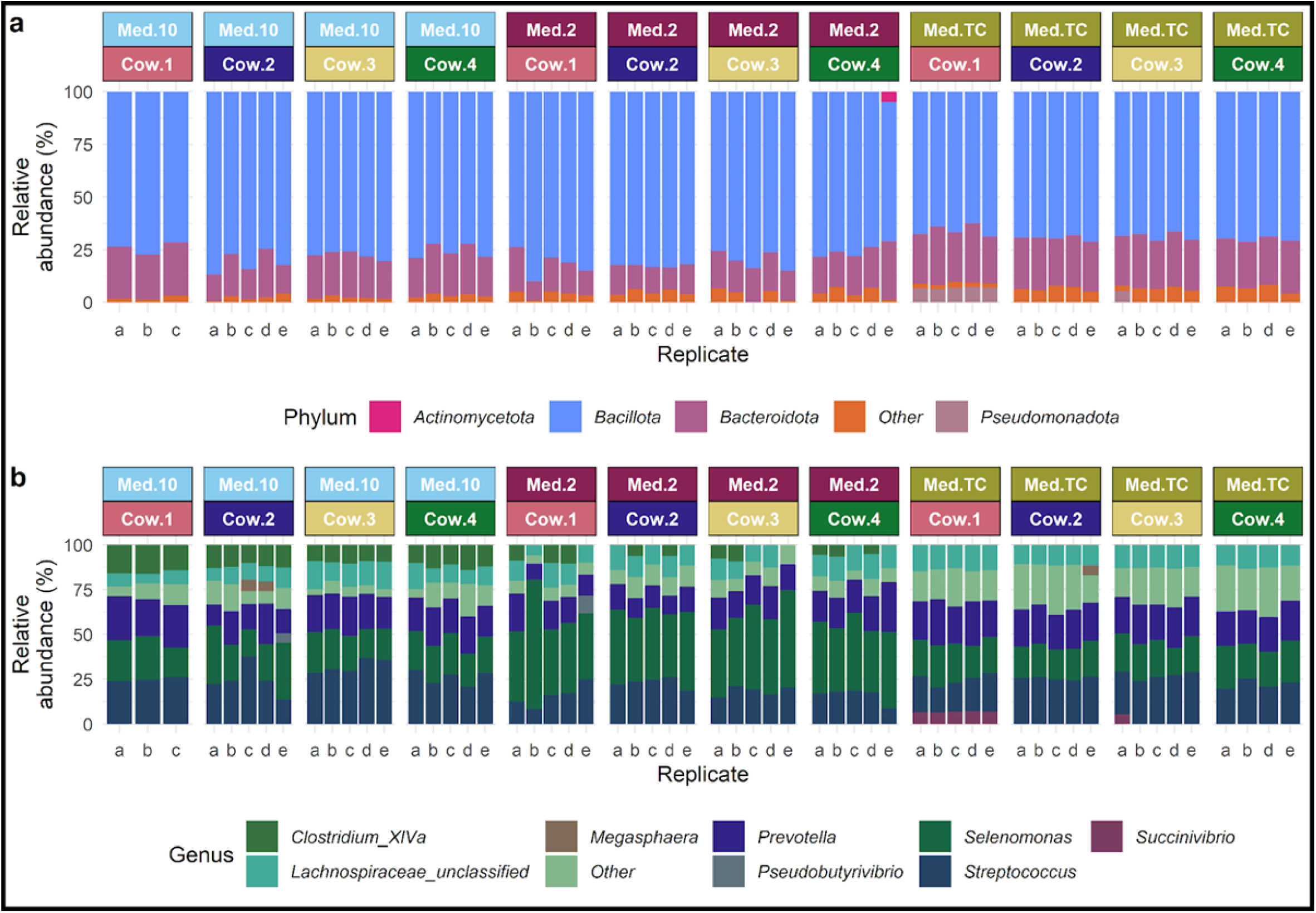
The relative abundances of phyla (a) and genera (b) found in each replicate culture by cow and medium. Any phyla or genera with a relative abundance < 5% are included in the ‘Other’ category.

The α-diversity, or species richness, of a community is defined as the variation of taxa within a community, and can be measured in a multitude of ways (29). Common indices of α-diversity include the Observed diversity which takes into account only the number of taxa and the number of individuals, Chao1 diversity which also includes the number of singletons and doubletons in that community to calculate species richness, and Shannon diversity which takes into account the relative and absolute abundance of different taxa combining both the diversity and evenness of the community (29,30). When comparing the media Med2 and MedTC, there was a significant difference between the α-diversity for all three indices using a one-way ANOVA and post-hoc Tukey HSD test (Observed: p = 3.48E-03, 95% C.I. = 1.58,9.18; Shannon: p = < 1.00E-06, 95% C.I. = 0.52,0.79; Chao1: p = 0.03, 95% C.I. = 0.27,7.78) (Figure 2). For the other media comparisons, there was no significant difference between the Observed and Chao1 diversities, however there was a significant difference between the Shannon diversity between the media Med2 and Med10 (p < 1.00E-06, 95% C.I. = 0.53,-0.26) and Med10 and MedTC (p = 1.29E-04, 95% C.I. = 0.12,0.39). When looking at the difference in α-diversity metrics between the microbial communities grown from different individual cows, irrespective of media, there were no significant differences between the Observed, Shannon, or Chao1 diversities.

**Figure 2.**
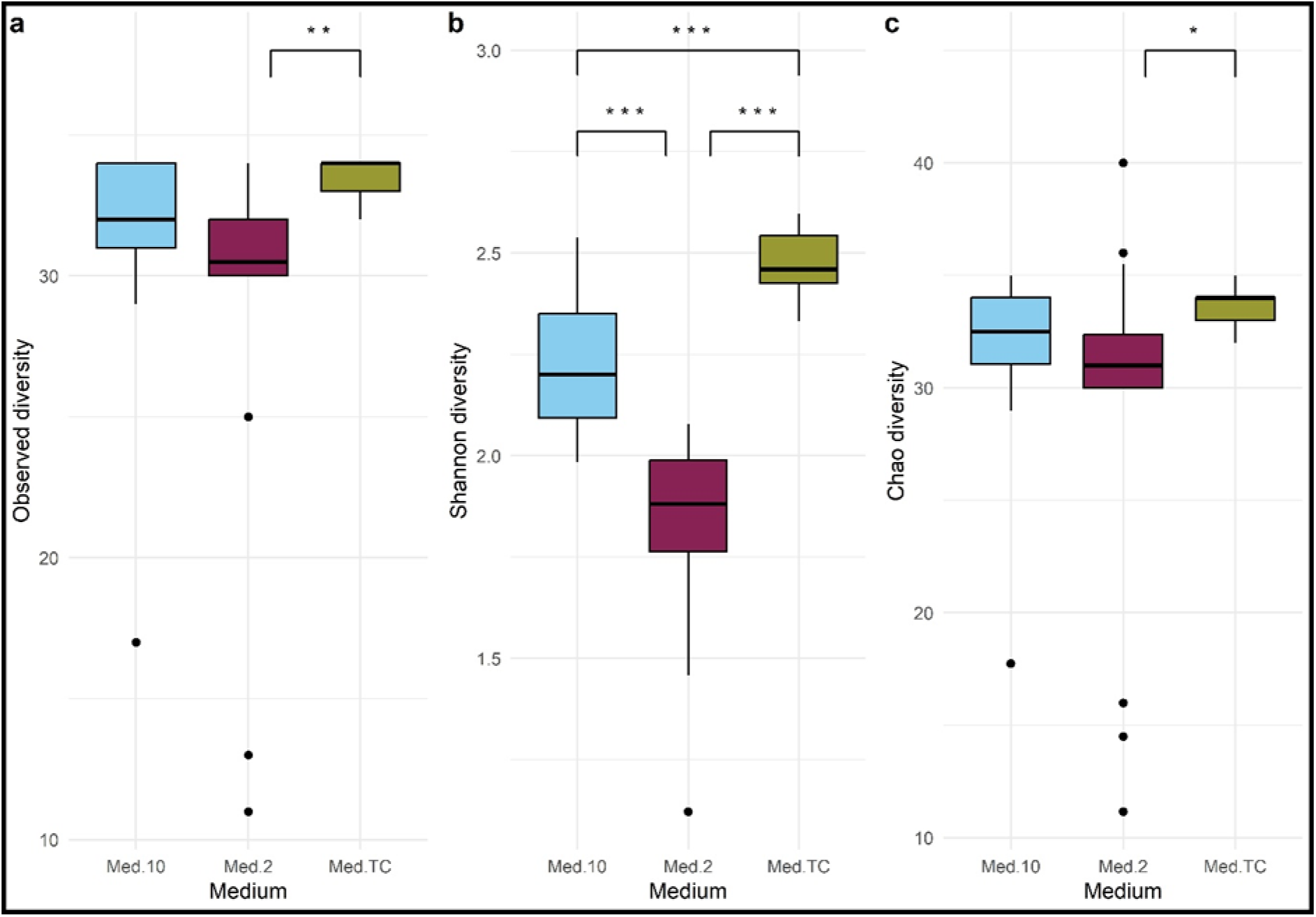
Comparison of α-diversity metrics between the microbial communities grown on the three media. a) Observed diversity, b) Shannon diversity, c) Chao1 diversity. Statistical tests used are one-way ANOVAs followed by post-hoc Tukey HSD tests if p < 0.05, with the results indicated with P < 0.05 = *, P < 0.01 = **, P < 0.001 = ***.

β-diversity is another method used to measure the diversity variation between communities 22), with the UniFrac metric along with Principal Coordinates Analysis (PCoA) often used to investigate differences between microbial populations, taking into account the phylogeny of the microbial taxa (31). There was a significant difference in the β-diversity in the microbial communities found in the different media (PERMANOVA, F(2, 54, 56) = 2.67, p = 0.03), though post-hoc pairwise comparisons using a pairwise-PERMANOVA with Bonferroni correction for multiple testing only indicated a significant difference between the media Med2 and MedTC (F(1, 37, 38) = 4.11, p = 0.001) (Figure 3a). Using the same tests, there was no significant difference in the β-diversity between samples from different cattle (Figure 3b).

**Figure 3.**
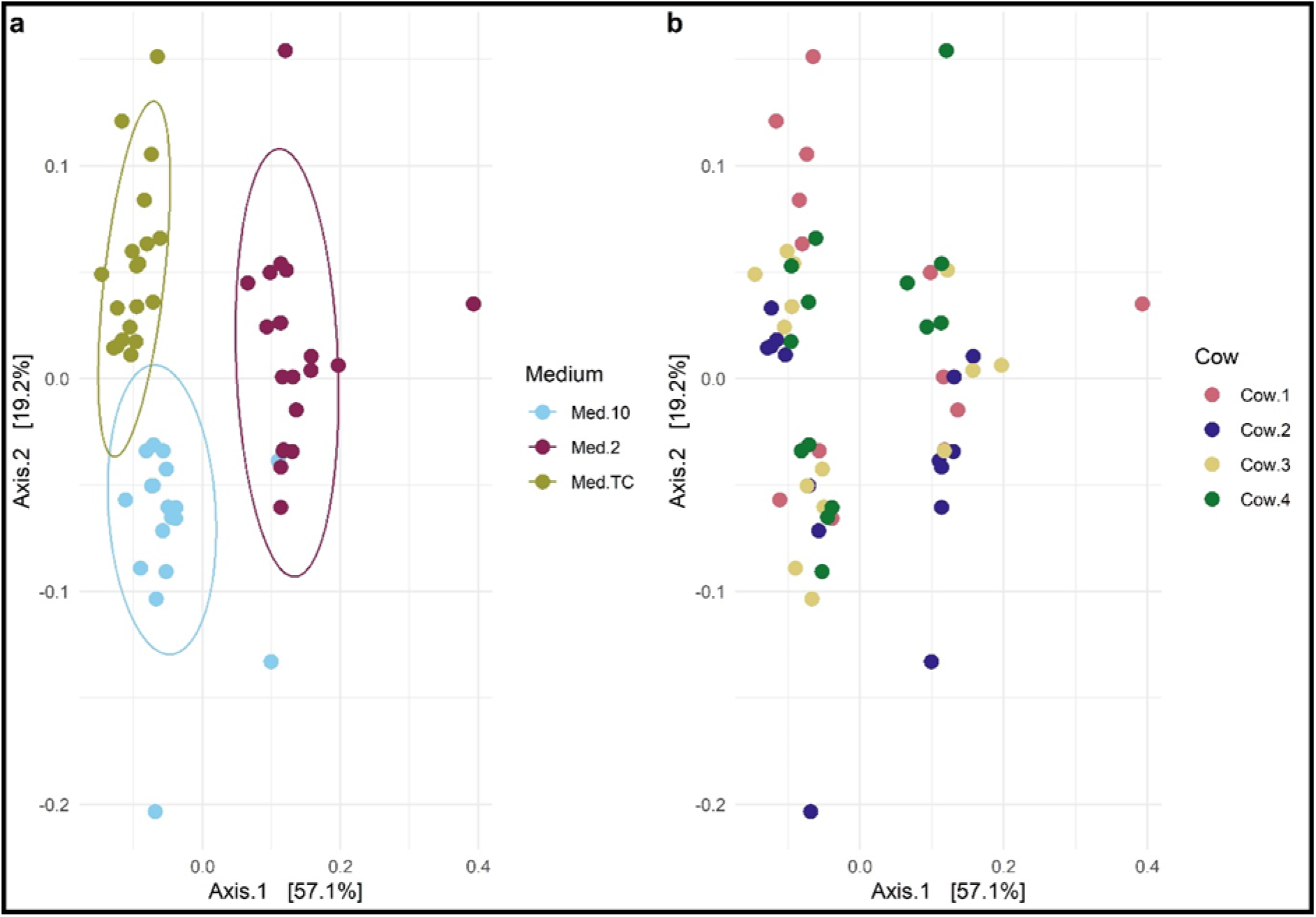
PCoA plot of weighted UniFrac distances of the cultures highlighted by medium (a) and source cow (b). Statistical tests used are one-way PERMANOVAs followed by post-hoc pairwise-PERMANOVAs with Bonferroni correction tests if p < 0.05, with the results indicated with P < 0.05 = *, P < 0.01 = **, P < 0.001 = ***.

Of the 34 OTUs classified to genus level found in the cultures, 31 were found in a significantly different mean relative abundance in at least one medium compared to the others (Kruskal-Wallis rank sum test with Bonferroni correction for multiple comparisons, Supplementary Table 1). The remaining three were not found to be enriched in any of the media. For those OTUs found in a significantly different relative abundance by medium, a post-hoc Dunn test was carried out (Supplementary Table 2). The 8 most abundant OTUs found across all the cultured samples are shown in Figure 4, which, when combined, make up 80% of the OTUs in terms of relative abundance.

**Figure 4.**
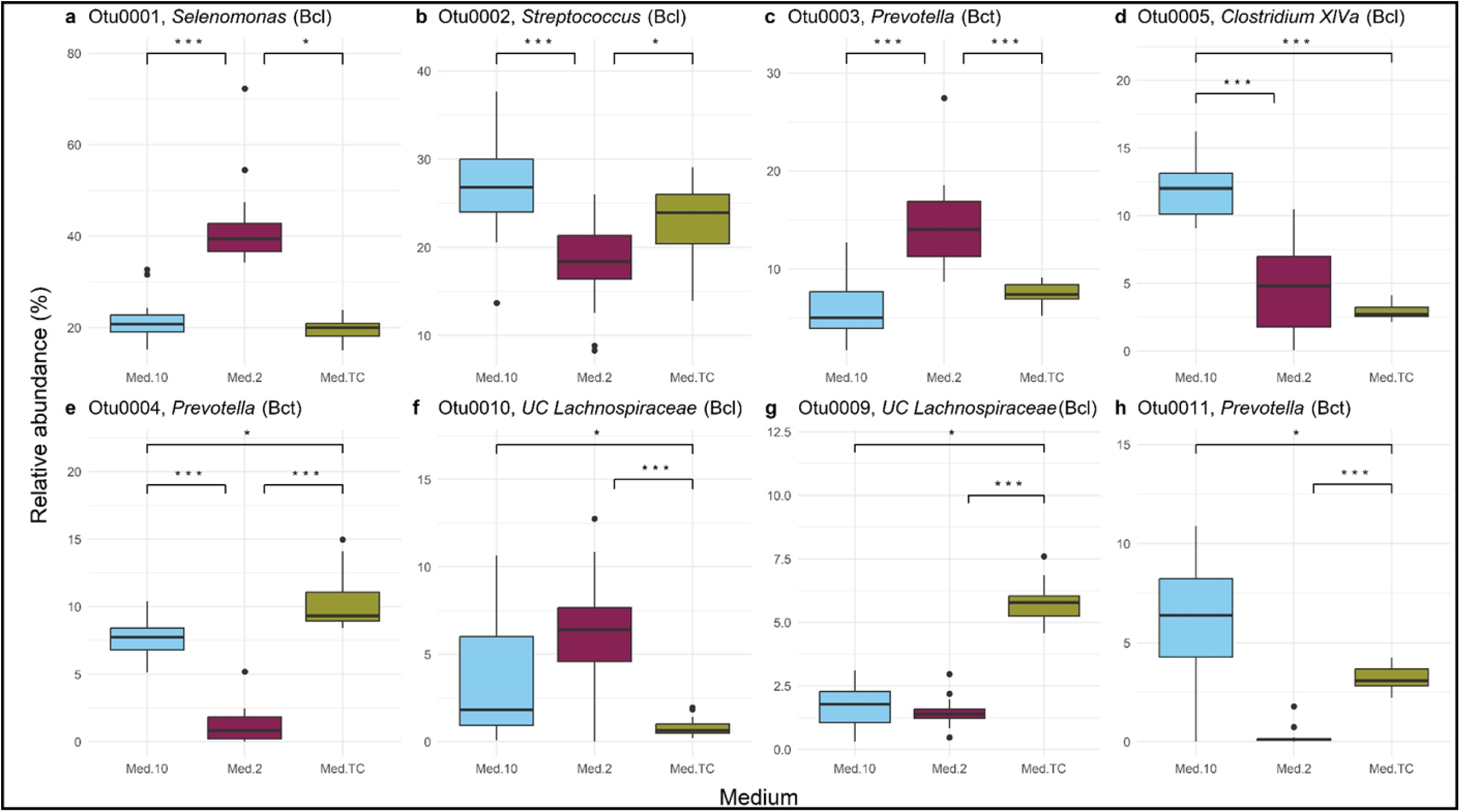
The eight most abundant OTUs (classified to genus level) and their differential relative abundances in the cultured samples by medium. The taxa included are; a) Selenomonas, b) Streptococcus, c) Prevotella, d) Clostridium XIVa, e) Prevotella, f) unclassified Lachnospiraceae, g) unclassified Lachnospiraceae, h) Prevotella. Genera from the Bacillota phylum are indicated with ‘Bcl’, and those from the Bacteroidota phylum are indicated with ‘Bct’. Statistical tests used are Kruskal-Wallis tests followed by post-hoc pairwise Dunn tests with Bonferroni correction if p < 0.05, with the results indicated with P < 0.05 = *, P < 0.01 = **, P < 0.001 = ***.

### 4.4 Microbes identified in the source rumen fluid inoculum

In addition to sequencing the 16S V4 region found in the cultures, the rumen fluid from each cow used as the culture inoculum was also sequenced. After going through the same filtering steps as the cultures, the four samples of rumen fluid contained 34 OTUs (classified to genus level). Of these, *Bacteroidota* (52.53% ± 5.10) was the most common phylum found in the four cows rumen fluid, based on their relative abundance, followed by *Bacillota* (41.00% ± 3.96), the archaeal phylum *Euryarchaeota* (5.12% ± 1.94), *Pseudomonadota* (1.22% ± 0.78), and finally *Actinomycetota* (0.12% ± 0.08) (Figure 5a). The most common genera found in the four cows rumen fluid, based on their relative abundance, were *Prevotella* (29.13% ± 4.16), *Butyrivibrio* (18.21% ± 2.08), *Succiniclasticum* (15.57% ± 5.03), unclassified *Bacteroidetes* (13.91% ± 1.67), and unclassified *Prevotellaceae* (9.50% ± 2.01) (Figure 5b).

**Figure 5.**
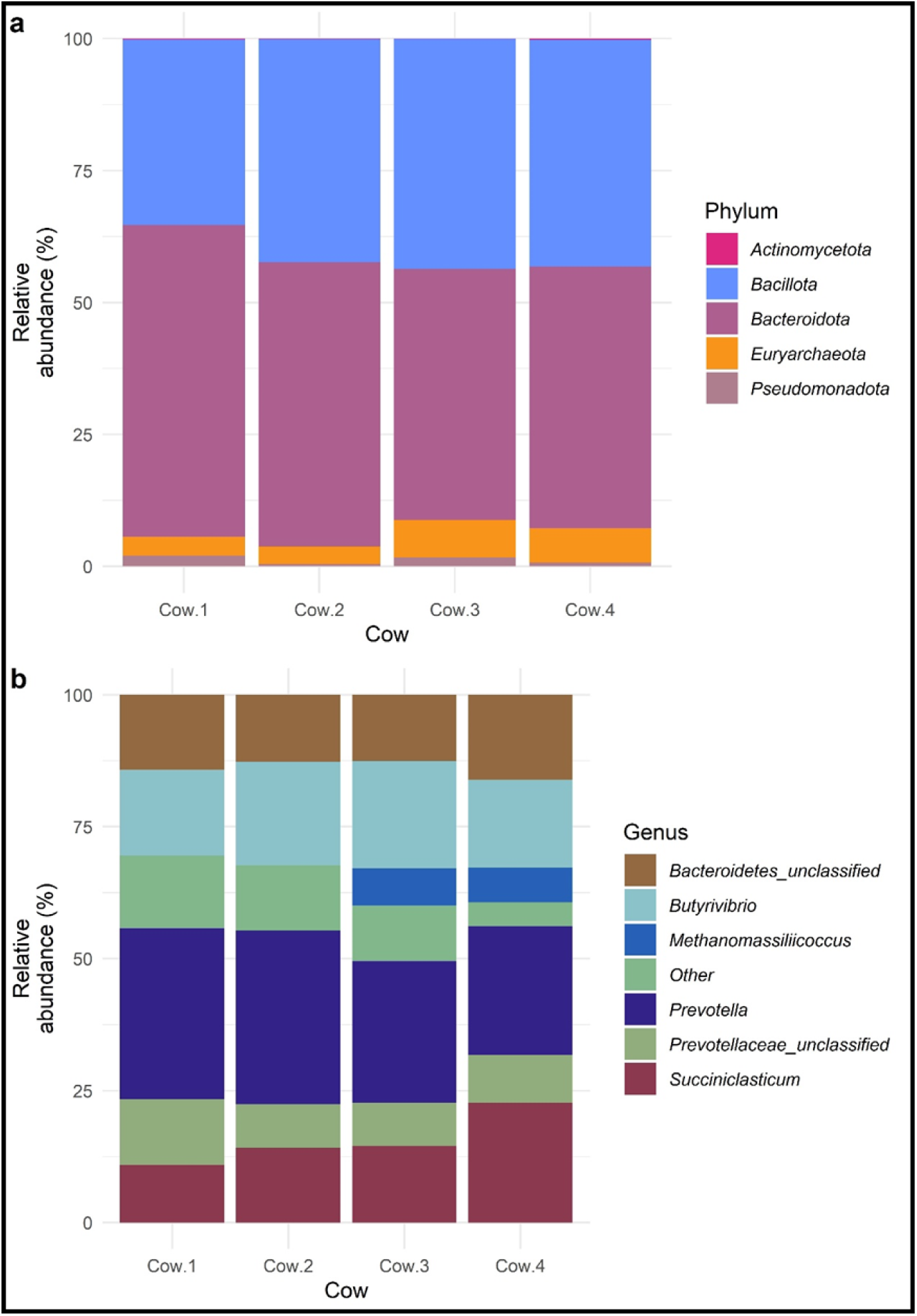
The relative abundances of the phyla (a) and genera (b) that make up the microbial communities found in each of the rumen fluid samples from the source cows. All phyla are shown, however any genera with a relative abundance < 5% are included in the ‘Other category.

### 4.5 The difference between the cultured microbes and those in the source rumen fluid inoculum

When comparing the microbial communities found in the rumen fluid from the source cows to those grown in culture, vast differences were observed (Figure 6, Supplementary Table 3). The same OTUs were found in both datasets, however in different relative abundances. This was quantified using the Log2-fold change of the mean relative abundance in the different media compared to the rumen fluid. Based on this, the phyla (Figure 6a) *Actinomycetota* (Log2-fold change compared to rumen fluid; Med10 = 3.66, Med2 = 4.60, MedTC = 3.84) and *Bacillota* (Med10 = 0.92, Med2 = 0.96, MedTC = 0.74) were overrepresented in the cultures on all media, and *Pseudomonadota* was overrepresented in Med2 (0.29) and MedTC (2.02), but not Med10 (−0.40). *Euryarchaeota* archaea (Med10 = −6.93, Med2 = −9.92, MedTC = −3.32) and *Bacteroidota* (Med10 = −1.39, Med2 = −1.73, MedTC = −1.11) were found in lower relative abundances in the media compared to the rumen fluid. Regarding genera (Figure 6b), the most underrepresented in the media compared to the rumen fluid were the methanogenic *Methanomassiliicoccus* in Med2 (−9.92), *Succiniclasticum* in Med2 (−8.47), and unclassified *Prevotellaceae* in Med2 (−8.26). The genera most overrepresented in culture were *Clostridium XlVa* (Med10 = 11.23, Med2 = 9.91, MedTC = 9.17), *Megasphaera* (Med10 = 8.34, MedTC = 8.99), *Selenomonadales* in Med10 (7.16), and *Streptococcus* (Med10 = 7.87, Med2 = 7.31, MedTC = 7.64).

**Figure 6.**
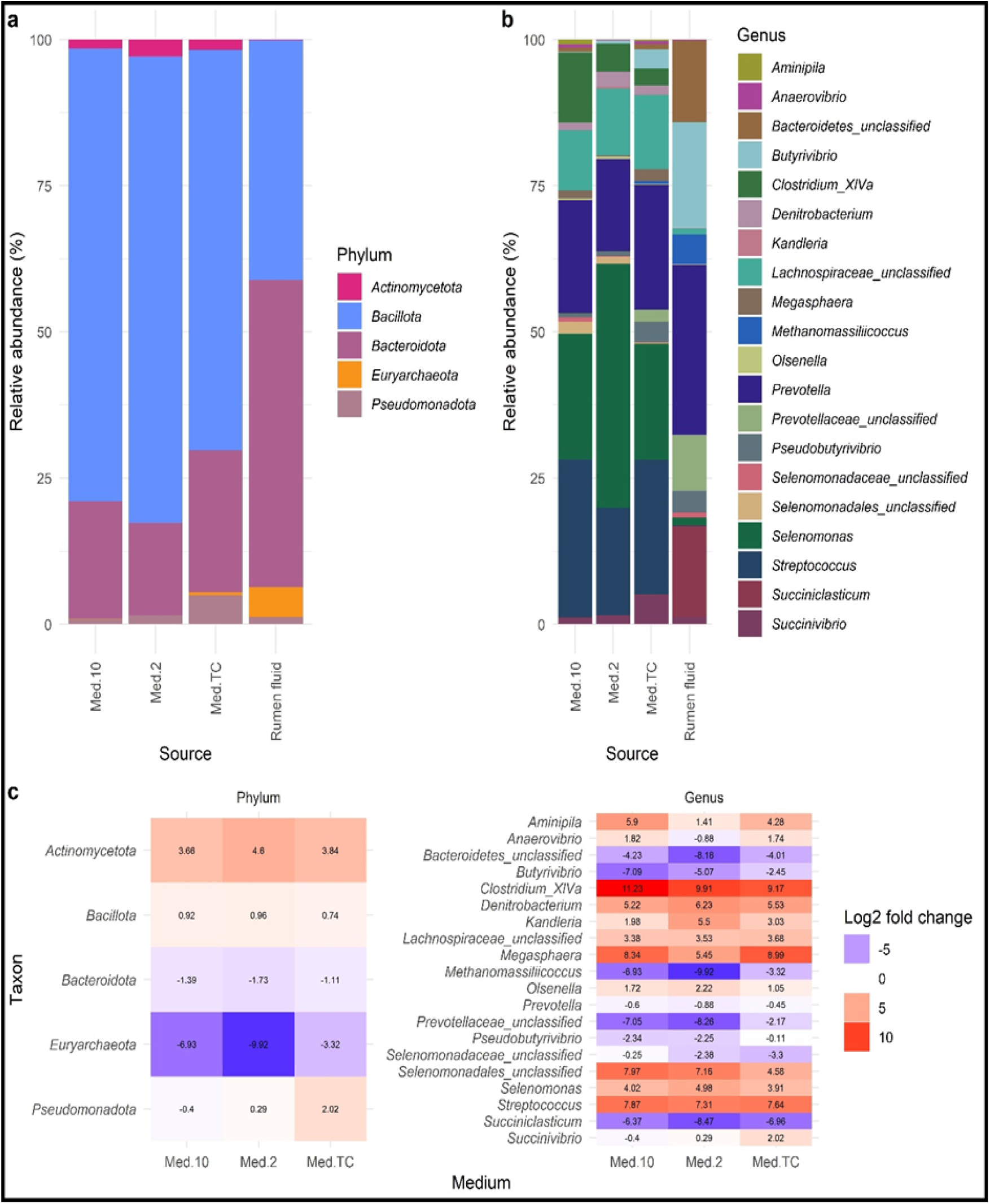
The mean relative abundances of the phyla (a) and genera (b) that make up the microbial communities found in the cultures grown on each of the types of media compared to the rumen fluid samples, along with the Log2 fold change of the mean relative abundance of each phylum and genus compared to the respective mean relative abundance found in the rumen fluid. Those taxa found in higher mean relative abundances in the rumen fluid are highlighted in red, those in lower mean relative abundances are shown in blue, and those with little change are shown in white.

## 5 Discussion

Culturing rumen microbes is essential to explore the full diversity of the rumen microbiome, and investigate how different microbes interact and react to different conditions. However the majority of rumen microbes are fastidious obligate anaerobes, making growing them in culture very difficult. In addition, many of the microbes in the rumen, identified through metagenomic methods, cannot be classified and have not been isolated in pure culture, meaning it is difficult to develop culture media that will support their growth. In this study, using a range of commonly used media to isolate rumen microorganisms, 34 rumen microbes were cultivated in co-culture. Different rumen microbial genera, such as *Selenomonas*, *Streptococcus*, and *Prevotella*, grew in different relative abundances on the different media, and a number of microbes were grown that could not be classified to genus level. This shows the value of different media for selecting for the growth of different rumen microbes, and the potential for further optimisation to enrich for the growth of target, as-yet-unclassified, microbes.

The majority of the OTUs identified in the cultures belonged to the genus *Selenomonas*, a Gram-negative, obligately anaerobic, flagellated genus of bacteria, with those found in, and isolated from, the rumen usually belonging to the species *S. ruminantium* (32,33). In this study, the single OTU identified as belonging to the genus *Selenomonas* grew in the highest relative abundance in the medium with no VFAs and the highest amount of yeast extract and also contained many less complex carbohydrates such as cellobiose, maltose, and glucose (Med2). This, along with the presence of sodium lactate (34,35), could explain why this microbe grew in the highest relative abundance on this medium. The next most abundant microbe cultured in this study were *Streptococcus spp*., a genus of Gram-positive facultative aerobes (36), likely to be from the bovis species group which includes, *S. bovis* (now *S. equinus*) and *S. gallolyticus* (36). They appear in the highest relative abundance on Med2 and MedTC, which may be due to the presence of sterile clarified rumen fluid, mineral solution 5, sodium lactate, NaHCO_3_, and lack of VFAs compared to Med10. *Prevotella*, another microbe cultured in high abundance, are rod-shaped and vary in length, being non-motile, non-sporing, and strictly anaerobic (37,38). To grow they require haemin, menadione, and peptides such as those found in trypticase, peptone, and yeast extract (38), at least one of which is found in all of the media used here. The three most relatively abundant OTUs belonging to *Prevotella* were enriched on one of each medium, and could be due to them belonging to different species or strains and therefore having different substrate requirements. Finally, the next most abundant microbe cultured was a form of unclassified *Lachnospiraceae*, potentially belonging to one of the existing genera, such as *Butyrivibrio*, *Lachnospira*, and *Roseburia*, or a new lineage entirely (33). These genera are all obligately anaerobic, motile, curved rods, with various Gram staining characteristics (32,33), and nutrient requirements (such as a wide variety of carbohydrate sources (33,39)) seemingly effectively met by all the media. The two most relatively abundant OTUs belonging to this unclassified taxon were also enriched in different media, suggesting they are different taxa with different metabolic requirements.

The complete bacterial and archaeal microbiomes of the cattle used in this study were similar, aligning with evidence suggesting that similarly treated animals of the same breed raised in the same location will have a similar microbiome (4). The majority of the OTUs identified in the full rumen fluid belonged to the phyla *Bacteroidota*, *Bacillota*, unclassified *Bacteria*, and the archaeal *Euryarchaeota*. This coincides with the generally agreed structure of the core microbiome outlined in reviews such as Terry et al. (2019) (40), Castillo-González et al. (2014) (41), and Anderson et al. (2021) (42). However the rumen microbiome structure between ruminant populations can differ due to the location of the animals (43) and their breed (44). Host diet also affects the rumen microbiome, with cattle used in this study grazed on perennial ryegrass with a supplementary garlic lick. A grass diet is often associated with higher levels of *Bacteroidota* (45), and garlic licks are associated with reduced levels of methanogens (46) and increased levels of *Prevotella* (47).

The majority of the rumen microbiome is uncultured and taxa were not able to be identified or classified using current databases (15–18,48,49). At the moment, isolates of *Bacillota*, *Pseudomonadota*, and *Actinomycetota* are found most often in culture collections with *Bacteroidota* being under represented (16). This is also observed in this study with a higher proportion of the bacteria cultured belonging to *Bacillota*, than those in the rumen fluid where a large proportion belong to *Bacteroidota.* Even when using a broad range of media that enrich for different microbes, and employing a dilution effect which should select for the most abundant rumen microbes, the profile of the microbes cultured is vastly different to that of those in the original rumen fluid.

Despite this the bacteria cultured here, and enriched in different media types, have been linked to key processes in the rumen, and the microbes cultured here coincide with others who have used Med10 (50,51), Med2 (52–56), and MedTC media (57). Higher abundances of *Selenomonas spp*. are associated with high grain (32,33) or high concentrate diets (58). Xue et al. (2022) found that more efficient animals showed interactions in the rumen between these *Selenomonas spp*. and bacteria of the *Succinivibrionaceae* family (59), with Smith et al. (2022) observing higher levels of *Selenomonas* in animals with low residual methane emissions (60). High levels of the lactate degrading species, often named *Selenomonas ruminantium subsp. Lactilytica*, are often found in higher numbers in the rumen when the rumen pH is reduced to the extent it leads to acidosis and bloat. As a genus involved in propionate production (34), further study of this genus in culture could lead to a better understanding of how this pathway could be used to reduce methane production. In addition, a probiotic dose of a strain of *Selenomonas ruminantium subsp. Lactilytica* was able to reduce the effects of lactic acidosis (61), and could act as a potential treatment or prophylactic for this issue. High levels of *Streptococcus spp*. in the rumen can lead to acidosis caused by the accumulation of lactate (62), though ruminal *Streptococcus spp*. have been shown to produce bacteriocins that are inhibitory to other *Streptococci* (63), which could provide scope for the control of ruminal acidosis using bacteriocins or bacteriocin-producing bacteria. *Streptococcal* species are also one of the causative agents of mastitis (64), another disease these potential bacteriocins could help treat (65). *Prevotella spp*. also have the potential to promote the stimulation of alternative hydrogen sinks and reduce the amount of methane produced from ruminal fermentation (66).

In the literature, high abundance of *Prevotella spp*. seems to be linked to low methane production due to the shift in fermentation profile towards propionate production (66,67). However, *Prevotella spp.* are greatly under-represented in pure culture collections (68). Similarly to *Selenomonas spp*., this may have the potential to improve ruminant efficiency if the proportion of *Prevotella spp*. are increased in the rumen, however in-depth study of different species and strains *in vitro* is required first. Many unclassified *Lachnospiraceae* have been identified in the rumen, and are likely to be important (48). *Lachnospiraceae* have been shown to be associated with a high residual feed intake (69) and low feed conversion ratio (70), so the *in vitro* study and further classification of these isolates would help elucidate the as-yet-undetermined characteristics of these microbes, with the media used in this research able to facilitate this.

There is vast potential for the development of new innovative techniques and media formulations to culture and classify rumen microbes. These can include changes in sample collection, processing, and inoculation methods with innovations around media composition, incubation conditions, length of time incubated, and the type of anaerobic atmosphere needed. Though care has been taken to avoid as many caveats as possible, there are some confounding variables and biases in this research. For example, there are undoubtedly some dilution effects, with the 10^-2^ dilution of rumen fluid likely inhibiting the culture of rarer, less abundant microbes, and the culture techniques used here better targeting bacteria than archaea. The number of replicates and the incubation period length used were chosen as a balance of human and consumable resource management, while aiming to maximise the number of bacterial taxa captured in this experiment. This means that, similar to the previous point, less abundant taxa are likely to be excluded, along with those that are slow-growing. In addition, only one breed of cattle was sampled from, on one type of diet, meaning that microbes that might be more abundant in different breeds or diets will have been missed. There is also some current bias and accuracy of database choice on OTU classification (71), and the choice of the V4 16S region will also have meant that microbes such as fungi will have been missed in this study, and may have skewed some of the taxonomic classification (72). In addition, the difference in absolute amounts of these microbe between the three media could only be ascertained with additional qPCR.

There is great scope for expanding the number of cultured rumen microbes using different media and anaerobic techniques, as demonstrated in this study. Increasing numbers of these isolates in pure culture, and their availability in culture collections, is essential to improve knowledge surrounding the intricacies of the rumen microbiome, microbial degradation of feed products, and reduction of methane production. To work towards increasing the number of rumen microbes in culture collections and improving the understanding of the rumen microbiome, future work with these cultures will include using dilution-to-extinction and streak plating in order to isolate the bacteria in pure culture and to characterise these.

## 6 Author statements

## 6.1 Author contributions

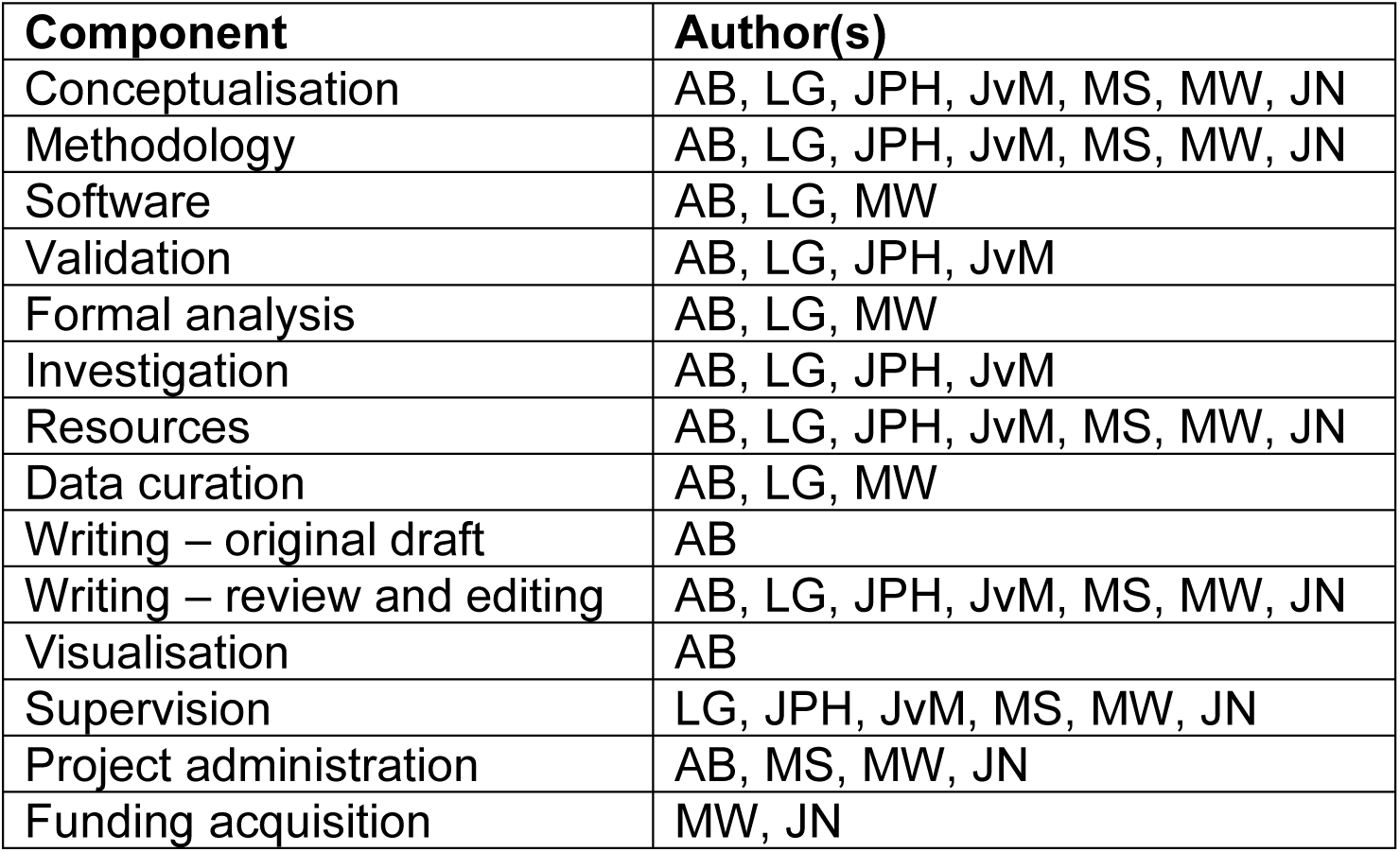

## 6.2 Conflicts of interest

The authors declare that there are no conflicts of interest.

## 6.3 Funding information

This work was supported by the Biotechnology and Biological Sciences Research Council (BBSRC) grant number BB/T00875X/1 and Institute Strategic Programme Grant BBS/E/RL/230001A. Sequencing was carried out by Edinburgh Genomics, the University of Edinburgh, which is partly supported with core funding from NERC (UKSBS PR18037). For the purpose of open access, the author has applied a CC-BY public copyright licence to any Author Accepted Manuscript version arising from this submission.

## 6.4 Ethical approval

Animal studies were carried out under, and adhered to, the guidelines and regulations of the UK Home Office Animals (Scientific Procedures) Act of 1986. The maintenance, sampling, and testing regime of these animals were covered by the Home Office project licence PP7153972. All experimental protocols were approved by the project licence holder and the University of Glasgow Animal Welfare and Ethical Review Board and adhere to ARRIVE guidelines.

## 6.5 Acknowledgements

The authors would like to thank the farm staff for facilitating sample collection.

## Notes

https://www.ebi.ac.uk/ena/browser/view/PRJEB78542

### Competing Interest Statement

The authors have declared no competing interest.

### Summary of Updates

The article has been reframed to focus more on the taxa grown on the media and how this compares to the microbial community found in the rumen rather than how much of the rumen microbiota can be cultured. No methods, analyses, or data have been changed. The title however has been completely changed. In addition the abstract, introduction, and discussion have been redrafted (keeping the majority of the information presented the same) to better fit the new focus and new title.

