## Supplementary Note 1 for "The selective culture and enrichment of major rumen bacteria on three distinct anaerobic culture media"

**Supplementary Note 1 Optimised protocol using the DNeasy® PowerLyzer® PowerSoil® Kit and QIAamp® Fast DNA Stool Mini Kits with additional modifications and steps used in this study.** In short, the QIAamp Fast DNA Stool Mini Kit protocol was followed with an additional glass bead beating step on centrifuged culture samples with the supernatant removed as described in the DNeasy® PowerLyzer® PowerSoil® Kit, and eluting the DNA in a lower volume.

1. Remove the 15ml centrifuge tubes containing the culture/sample pellet from the -80°C freezer and place tube on ice
2. Add 1 ml InhibitEX Buffer to each sample (QIAamp® Fast DNA Stool Mini Kit)
3. Vortex continuously for 1 min or until the stool sample is thoroughly homogenized (may need an extra 30 seconds to make sure all solids are re-suspended) (QIAamp® Fast DNA Stool Mini Kit)
4. Transfer 1ml of the solution to a 2ml Eppendorf
5. Incubate for 10 minutes at 70C (QIAamp® Fast DNA Stool Mini Kit – extended incubation)
6. Incubate at 5 minutes at 95C (Additional incubation step at higher temperature)
7. Vortex for 15 seconds (QIAamp® Fast DNA Stool Mini Kit)
8. Transfer 1ml of the sample to a glass bead beating tube
9. Vortex briefly
10. Lyse the sample using the FastPrep 24 for 45 seconds at 4.5m/s (DNeasy® PowerLyzer® PowerSoil® Kit Handbook)
11. Centrifuge bead beating tubes at for 30 seconds at 10,000g (DNeasy® PowerLyzer® PowerSoil® Kit Handbook)
12. Pipet 25 µL proteinase K into a new 2 ml Eppendorf (QIAamp® Fast DNA Stool Mini Kit)
13. Pipet 500 µL supernatant from step 10 into the 2 ml Eppendorf containing proteinase (QIAamp® Fast DNA Stool Mini Kit)
14. Add 600 µL Buffer AL and immediately vortex for 15 seconds (QIAamp® Fast DNA Stool Mini Kit)
15. Incubate at 70°C for 10 min (QIAamp® Fast DNA Stool Mini Kit)
16. Add 600 µL of ethanol (96–100%) to the lysate, and mix by vortexing (QIAamp® Fast DNA Stool Mini Kit)
17. Centrifuge briefly to remove drops from the inside of the tube lid (QIAamp® Fast DNA Stool Mini Kit)
18. Carefully apply 600 µL lysate from step 16 to the QIAamp spin column (QIAamp® Fast DNA Stool Mini Kit)
19. Close the cap and centrifuge at full speed for 1 min (QIAamp® Fast DNA Stool Mini Kit)
20. Place the QIAamp spin column in a new 2 ml collection tube, and discard the tube containing the filtrate (QIAamp® Fast DNA Stool Mini Kit)
21. Repeat steps 18-20 until all of the lysate has been loaded on the column (if the lysate has not completely passed through the column after centrifugation, centrifuge again until the QIAamp spin column is empty) (QIAamp® Fast DNA Stool Mini Kit)
22. Carefully open the QIAamp spin column and add 500 µL Buffer AW1 (QIAamp® Fast DNA Stool Mini Kit)
23. Centrifuge at full speed for 1 min (QIAamp® Fast DNA Stool Mini Kit)
24. Place the QIAamp spin column in a new 2 ml collection tube, and discard the collection tube containing the filtrate (QIAamp® Fast DNA Stool Mini Kit)
25. Carefully open the QIAamp spin column and add 500 µL Buffer AW2 (QIAamp® Fast DNA Stool Mini Kit)
26. Centrifuge at full speed for 3 min (QIAamp® Fast DNA Stool Mini Kit)
27. Discard the collection tube containing the filtrate (QIAamp® Fast DNA Stool Mini Kit)
28. Place the QIAamp spin column in a new 2 ml collection tube and discard the old collection tube with the filtrate (QIAamp® Fast DNA Stool Mini Kit)
29. Centrifuge at full speed for 3 min (QIAamp® Fast DNA Stool Mini Kit)
30. Transfer the QIAamp spin column into a new, labelled 1.5 ml Eppendorf and pipet 50 µL Buffer ATE directly onto the QIAamp membrane (QIAamp® Fast DNA Stool Mini Kit – lower elution volume)
31. Incubate for 5 min at room temperature (QIAamp® Fast DNA Stool Mini Kit)
32. Centrifuge at full speed for 1 min to elute DNA (QIAamp® Fast DNA Stool Mini Kit)
