## Supplementary figures and images for "The selective culture and enrichment of major rumen bacteria on three distinct anaerobic culture media"

### Supplementary Figure 1

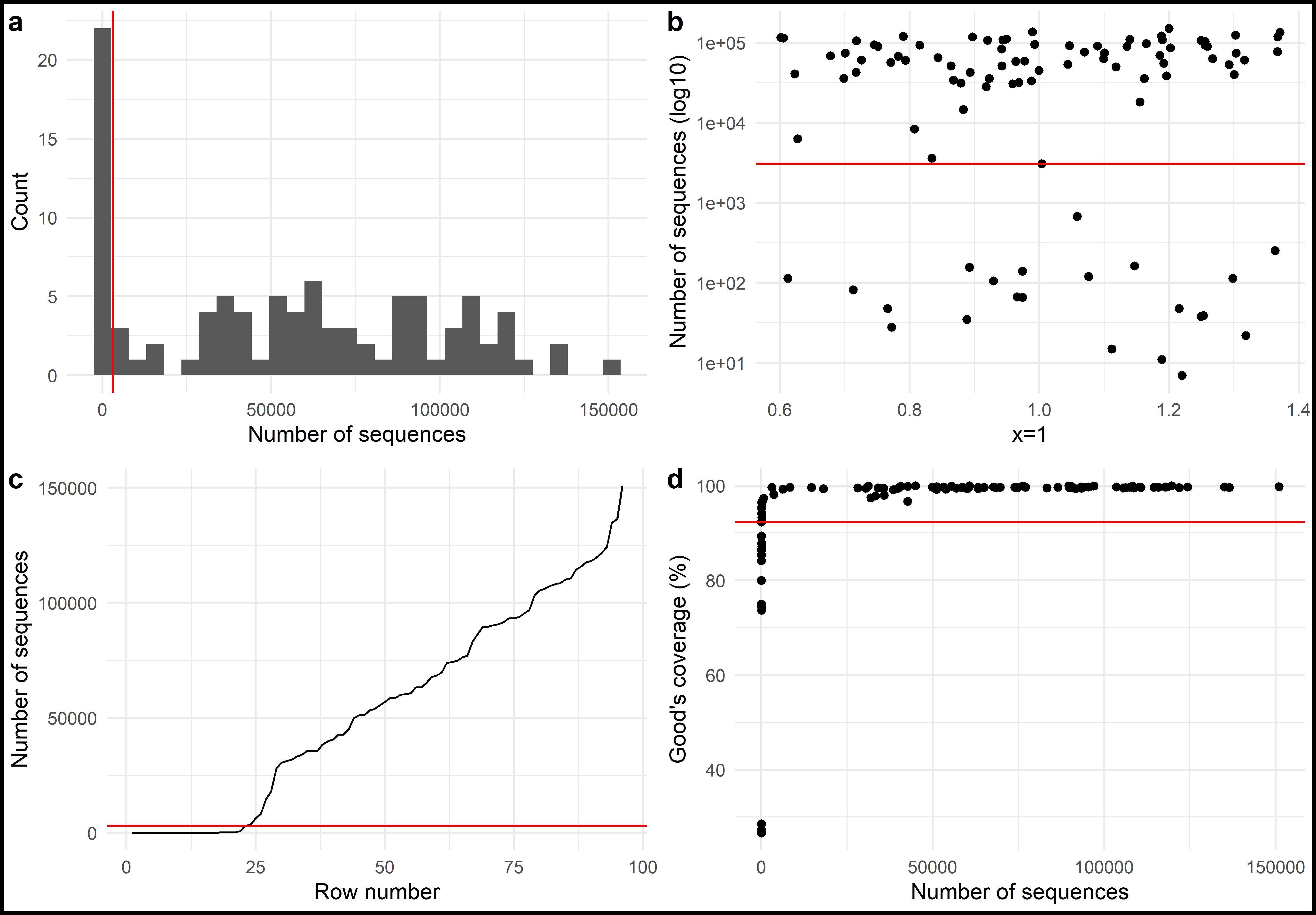

### Supplementary Figure 2

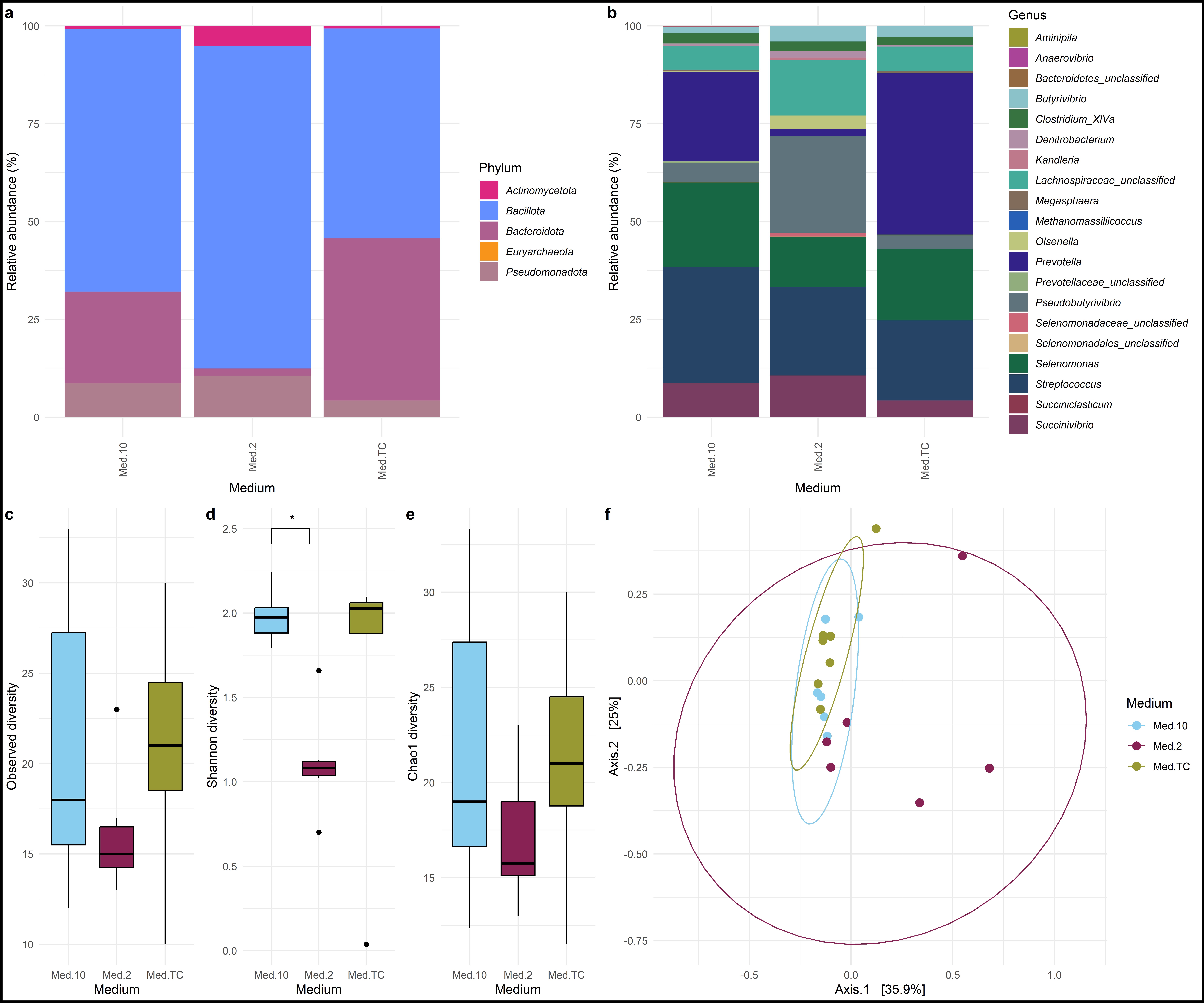

### Supplementary Figure 3

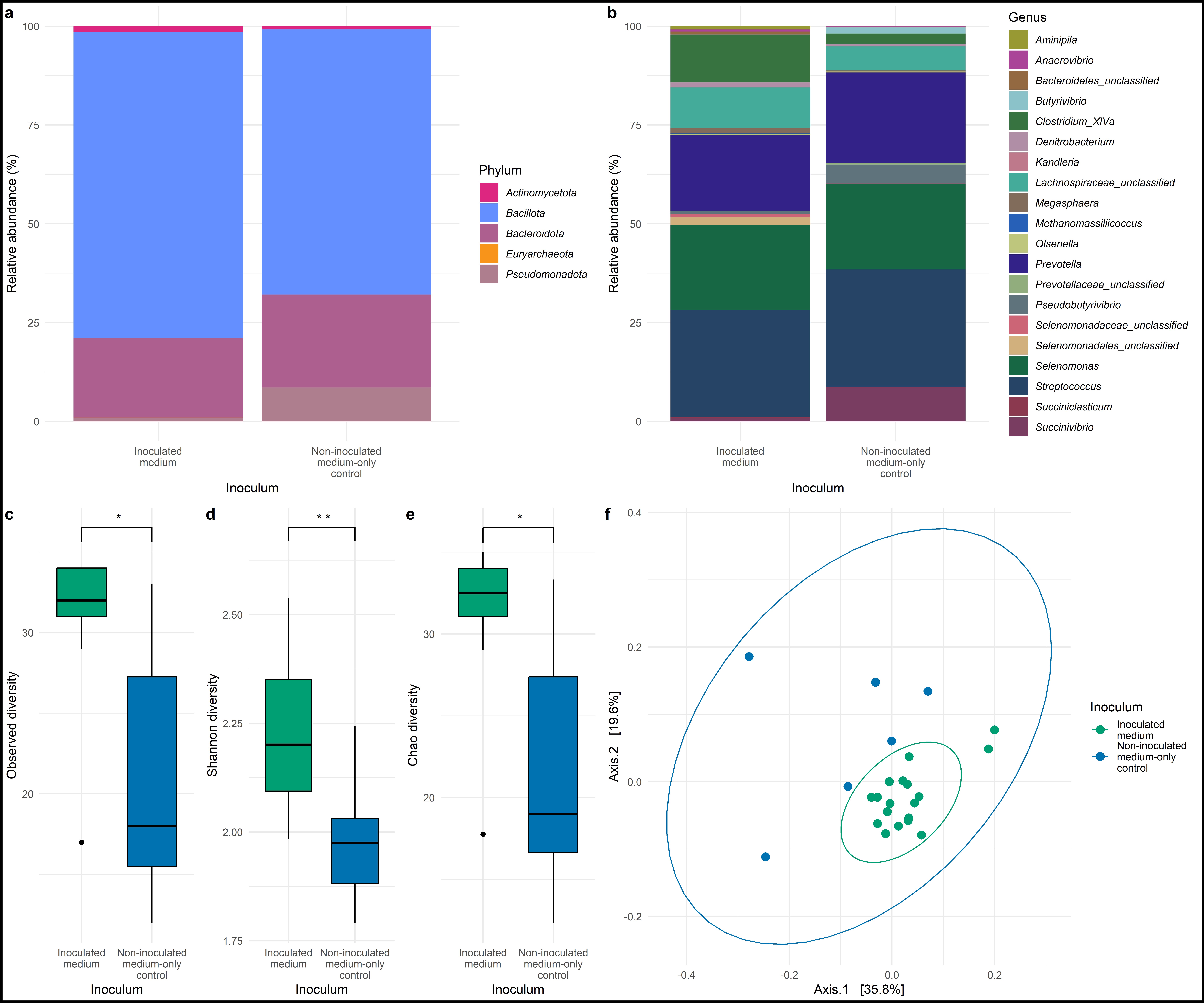

### Supplementary Figure 4

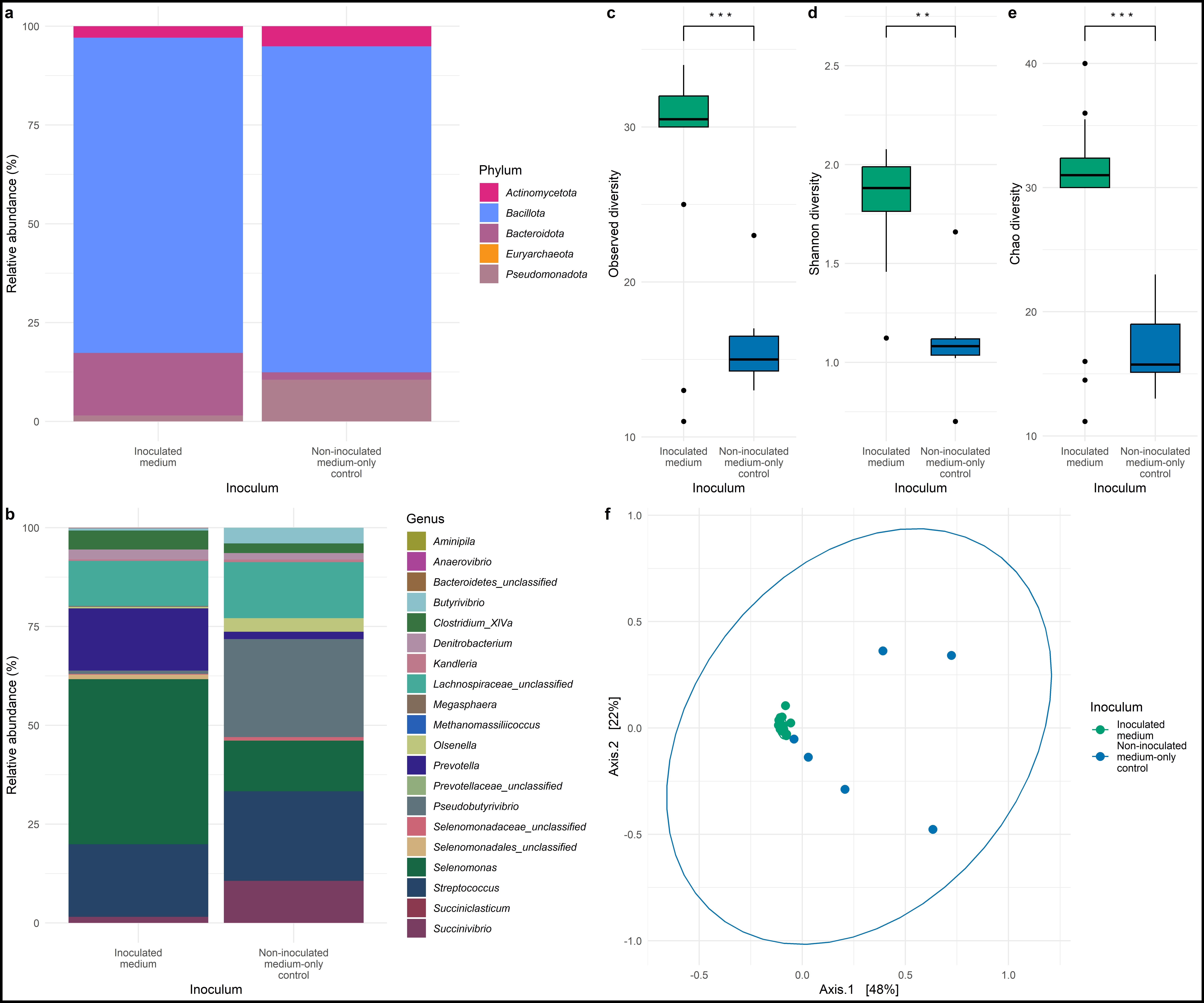

### Supplementary Figure 5

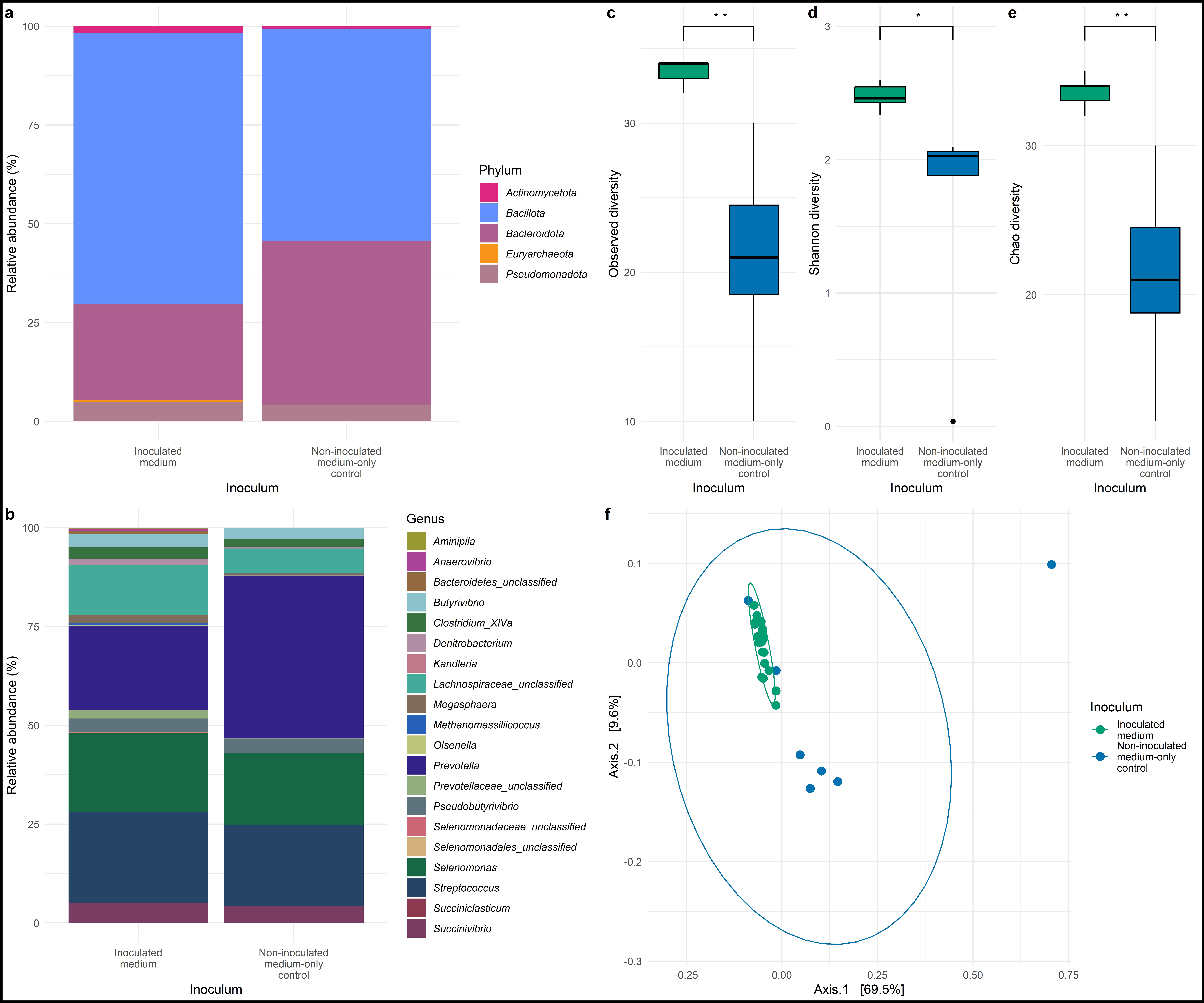

### Supplementary Figure 6

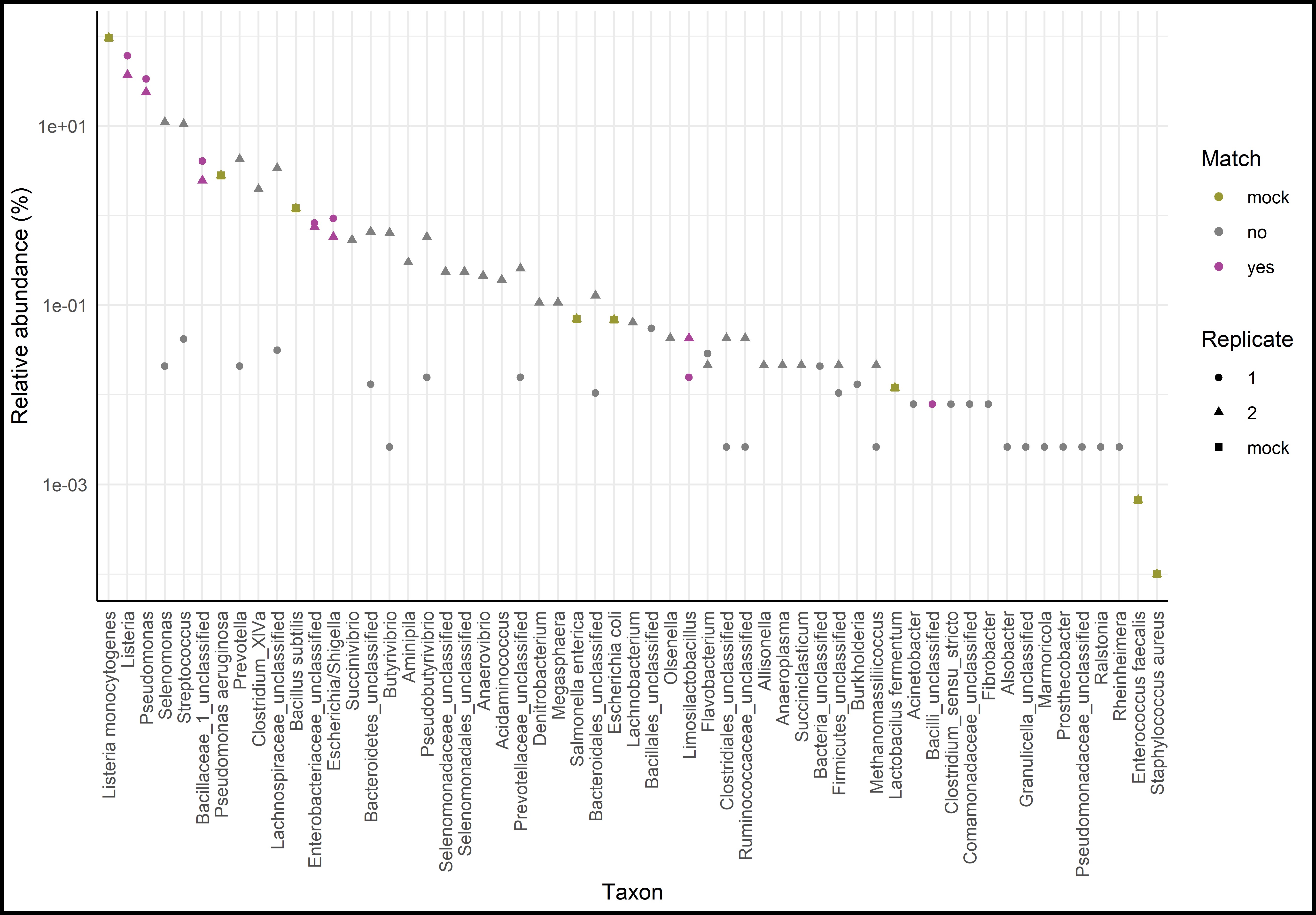
